# Sex and *APOE* Genotype Influence Respiratory Function Under Hypoxic and Hypoxic-Hypercapnic Conditions

**DOI:** 10.1101/2022.08.03.502692

**Authors:** Chase E. Taylor, Laura E. Mendenhall, Michael D. Sunshine, Jessica N. Wilson, Chris M. Calulot, Ramon C. Sun, Lance A. Johnson, Warren J. Alilain

**Affiliations:** Department of Neuroscience, University of Kentucky; Spinal Cord and Brain Injury Research Center, University of Kentucky; Department of Biochemistry & Molecular Biology, College of Medicine, University of Florida, Department of Biochemistry, University of Florida; Center for Advanced Spatial Biomolecule Research, University of Florida; Department of Physiology, University of Kentucky; Sanders-Brown Center on Aging, University of Kentucky

## Abstract

The apolipoprotein *(APOE)* gene has been studied due to its influence on Alzheimer’s disease (AD) development and work in an *APOE* mouse model recently demonstrated impaired respiratory motor plasticity following spinal cord injury (SCI). Individuals with AD often co-present with obstructive sleep apnea (OSA) characterized by cessations in breathing during sleep. Despite the prominence of *APOE* genotype and sex as factors in AD progression, little is known about the impact of these variables on respiratory control. Ventilation is tightly regulated across many systems, with respiratory rhythm formation occurring in the brainstem but modulated in response to chemoreception. Alterations within these modulatory systems may result in disruptions of appropriate respiratory control and ultimately, disease. Using mice expressing two different humanized *APOE* alleles, we characterized how sex and the presence of *APOE3* or *APOE4* influences ventilation during baseline breathing (normoxia) and during respiratory challenge. We show that sex and *APOE* genotype influence breathing during hypoxic challenge, which may have clinical implications in the context of AD and OSA. Additionally, female mice, while responding robustly to hypoxia, were unable to recover to baseline respiratory levels, emphasizing sex differences in disordered breathing.

## INTRODUCTION

Apolipoprotein E (ApoE) has 3 distinct alleles: *APOE2, E3*, and *E4* each coding for a different lipid transport protein variant ^1^. The *APOE4* allele is a key genetic determinant for late-onset Alzheimer’s disease (AD); it is estimated that heterozygous carriage of the *APOE4* allele results in a 4-fold increased risk of developing AD, while homozygous carriers exhibit a 15 fold increase ^2^. Approximately one in five people are carriers of the *APOE4* allele. Critically, in the context of our work, the *E4* Alzheimer’s connection has been linked to the presentation of cognitive decline and inflammation.

Breathing, the rhythmic inspiratory and expiratory patterns that facilitate gas exchange in the lungs, is essential to life and conditions impacting respiratory function and neural respiratory control are critical in the context of aging related disease ^3,4,5-7^. Obstructive sleep apnea (OSA) is a disease state which is characterized by cessations in breathing leading to periods of hypoxia ^8^. Repeated exposure to hypoxia leads to neuroinflammation and cognitive decline which can magnify the impacts of AD ^9^.

Considering the importance of *APOE4* as the key genetic determinant for AD and hypoxia as a major driver of cognitive decline, it is critical to understand the impact and relationship of the *APOE4* allele on neural respiratory control and the body’s innate response to hypoxia, i.e. the hypoxic ventilatory response (HVR) ^10^. We tested the hypothesis that *APOE4* negatively impacts breathing, particularly under conditions that would induce a response to respiratory stress. To test this hypothesis, we utilized whole body plethysmography (WBP) to measure ventilation in a humanized *APOE* mouse model under normoxia (21% O_2_), and two different gas challenges (hypoxia alone and hypoxic hypercapnia).

## METHODS

### Animals

All experiments were approved by the Institutional Animal Care and Use Committee at the University of Kentucky. Mice expressing human *APOE* isoforms under control of the endogenous mouse *APOE* promotor (targeted replacement mice) were backcrossed for at least 10 generations to the C57BL/6 background ^11,12^. Mice were group-housed by sex and genotype on a 12/12 light/dark cycle and fed normal chow diet *ad libitum*. Mice in cohort 1 (hypoxic challenge; n=33) weighed 36.9-48.6 g with ages ranging from 367-441 days. Mice in cohort 2 (hypoxic-hypercapnic challenge; n=16) weighed 32.02-52.7 g with ages ranging from 472-477 days. The average age and weight by sex, genotype, and cohort are listed in Table 1.

**Table 1.** Number, average weight, and average age of mice used in hypoxia and hypoxic-hypercapnia experiments. g = grams; SD = standard deviation; d = days

| Cohort | Sex | Genotype | n | Average weight (g) (SD) | Average age (d) (SD) |
| --- | --- | --- | --- | --- | --- |
| Hypoxia challenge | Male | E3 | 9 | 48.8 (4.7) | 410 (16.6) |
|  |  | E4 | 8 | 45.5 (4.6) | 441 (65.3) |
|  | Female | E3 | 8 | 36.9 (7.5) | 367 (10.1) |
|  |  | E4 | 8 | 37.5 (5.3) | 377 (9.0) |
| Hypoxic-hypercapnia challenge | Male | E3 | 4 | 52.7 (2.8) | 473 (0) |
|  |  | E4 | 4 | 47.5 (6.0) | 472.5 (4.0) |
|  | Female | E3 | 3 | 51.2 (5.9) | 477 (0) |
|  |  | E4 | 5 | 32.02 (4.2) | 476 (3.8) |

### Hypoxia and hypoxic-hypercapnia experiments

Whole body plethysmography (WBP,1 liter per minute flow) (DSI, Buxco FinePointe) was used to measure the ventilatory response during hypoxic (11% O_2_ balance nitrogen) and hypoxic-hypercapnic (7% CO_2_, 10.5% O_2_, balance nitrogen) challenge in adult mice. We calculated respiratory rate (breaths/min), tidal volume (mL/breath/g), and minute ventilation (mL/min/g) from the WBP flow traces. Whole body plethysmography recordings lasted approximately 90 minutes and occurred during daylight hours between 0930 and 1600 with food and water restricted during the 90-minute measurement period. To ensure consistency across all measures, the WBP system was recalibrated prior to every trial. Immediately prior to baseline readings, a 40-minute period of in-chamber acclimation occurred to reduce the prevalence of non-ventilatory respiratory behaviors associated with exploring the chamber during the baseline period. Acclimation was followed by a 30-minute period of normoxia (21% O_2_, balance nitrogen), then 10 minutes of respiratory challenge concluding with a 5-or 10-minute period of normoxia. The challenge design can be seen in Figure 1a and the number of animals per group is shown in Table 1. Following trials, mice were returned to their home cage in the vivarium. WBP chambers were thoroughly cleaned and dried between each group.

**Figure 1.**
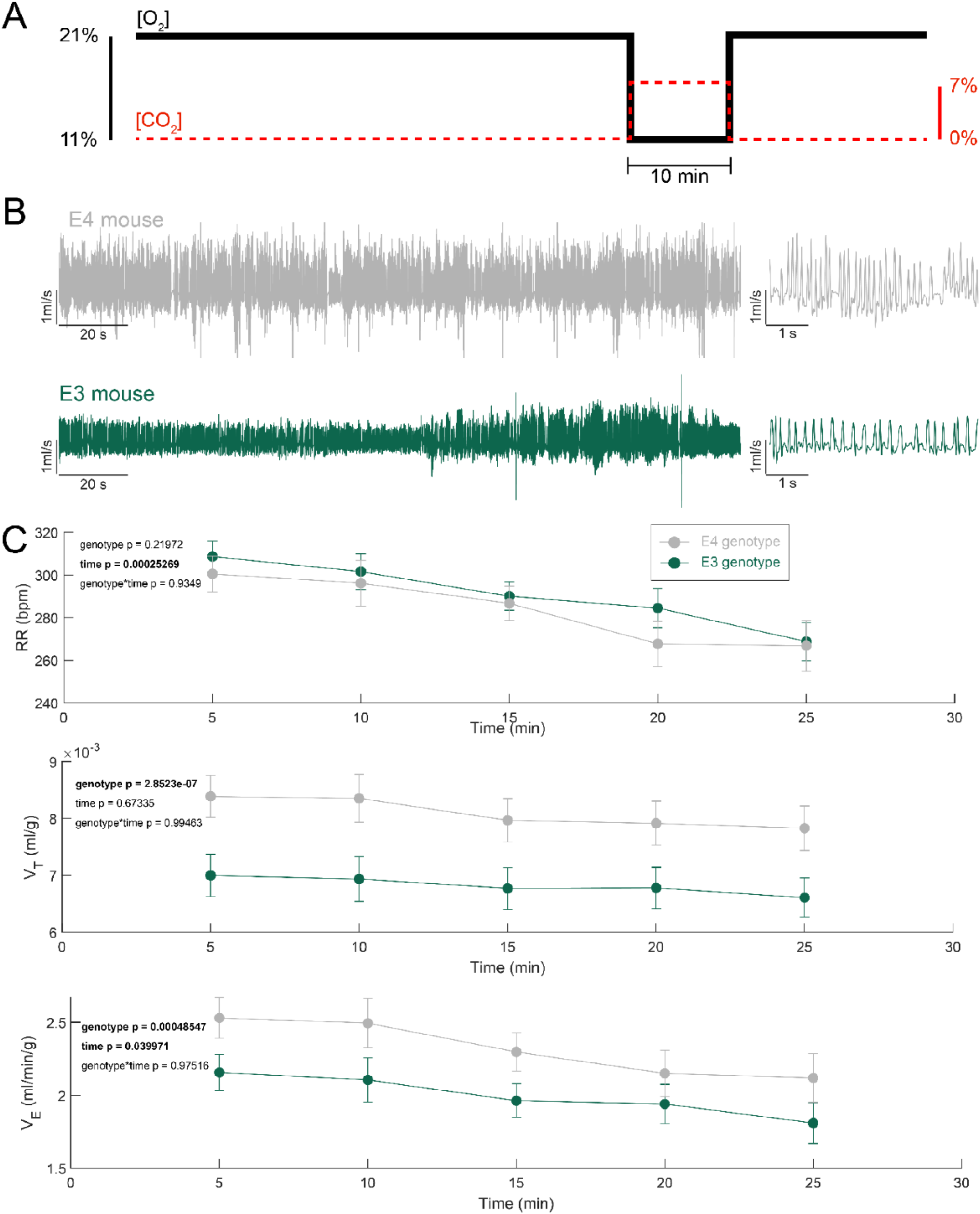
*APOE4* animals across sexes have increased tidal volume to maintain respiratory homeostasis under normoxic conditions while maintaining the same respiratory rate as *E3* mice. **a)** Timeline of WBP hypoxic and hypoxic hypercapnic challenges. **b)** Plethysmography airflow traces from a representative *APOE3* (green) or *APOE4* (gray) mouse during baseline (normoxia). **c)** Quantification of plethysmography during baseline (normoxia) in five-minute bins, testing for an overall effect of genotype prior to assessing sex differences (Two-way ANOVA mean ± SEM; n= 25 (*APOE4*, 13F/12M), n=24 (*APOE3*, 11F/13M)).

### Data analysis and statistical analysis

The normality and variance of the data was determined to ensure appropriate assumptions were met. If data within a particular experiment did not meet these assumptions non-parametric tests were used. Specific statistical tests for each figure are listed in the figure legends. Ventilatory measures were calculated in DSI FinePointe software and exported, breath by breath. The breath by breath data was used to calculate binned respiratory rate and average tidal volume, these metrics were multiplied to generate minute ventilation. This analysis and statistical testing was performed using MATLAB (MathWorks, Version 2021b). For hypoxic ventilatory challenge, hypoxic-hypercapnic ventilatory challenge, and post challenge measures, data were normalized to percent change relative from baseline. For statistical analysis, an ANOVA (or Kruskal-Wallis for non-parametric analyses) was used for statistical comparisons between sex and genotype. Tukey-Kramer post-hoc test was used to test for differences within sex or genotype. Results were considered statistically significant if p < 0.05. Investigators were not blinded to genotype or sex.

## RESULTS

### Both genotype and sex impact normoxic breathing with female mice showing the greatest tidal volume

To determine differences in baseline breathing, *APOE3* and *APOE4* male and female mice underwent a 25-minute period of normoxia prior to a 10-minute respiratory challenge. Representative flow traces from an *APOE3* and *E4* mouse (Figure 1B) illustrate larger tidal volume during baseline (normoxia) in the *APOE4* mouse. Quantification of these data showed an effect of genotype with *E4* mice having a larger tidal volume throughout the normoxic baseline (Figure 1C). Representative flow traces were shown from each sex and genotype (Figure 2A). Sex differences were present in tidal volume and minute ventilation when corrected for body weight (Figure 2B-D). Female mice had to breath at a deeper and greater weight corrected tidal volume relative to male mice to maintain baseline breathing (p<0.002, F_(1)_ = 11.96). Additionally, female mice had a larger weight corrected minute ventilation (p<0.03, F_(1)_ = 5.05). These baseline measures show a sex-dependent difference present that may affect how male and female mice will respond to a ventilatory challenge. Finally, *E4* mice had larger weight corrected tidal volume compared to *E3* mice (p<0.03, F_(1)_ = 5.50).

**Figure 2:**
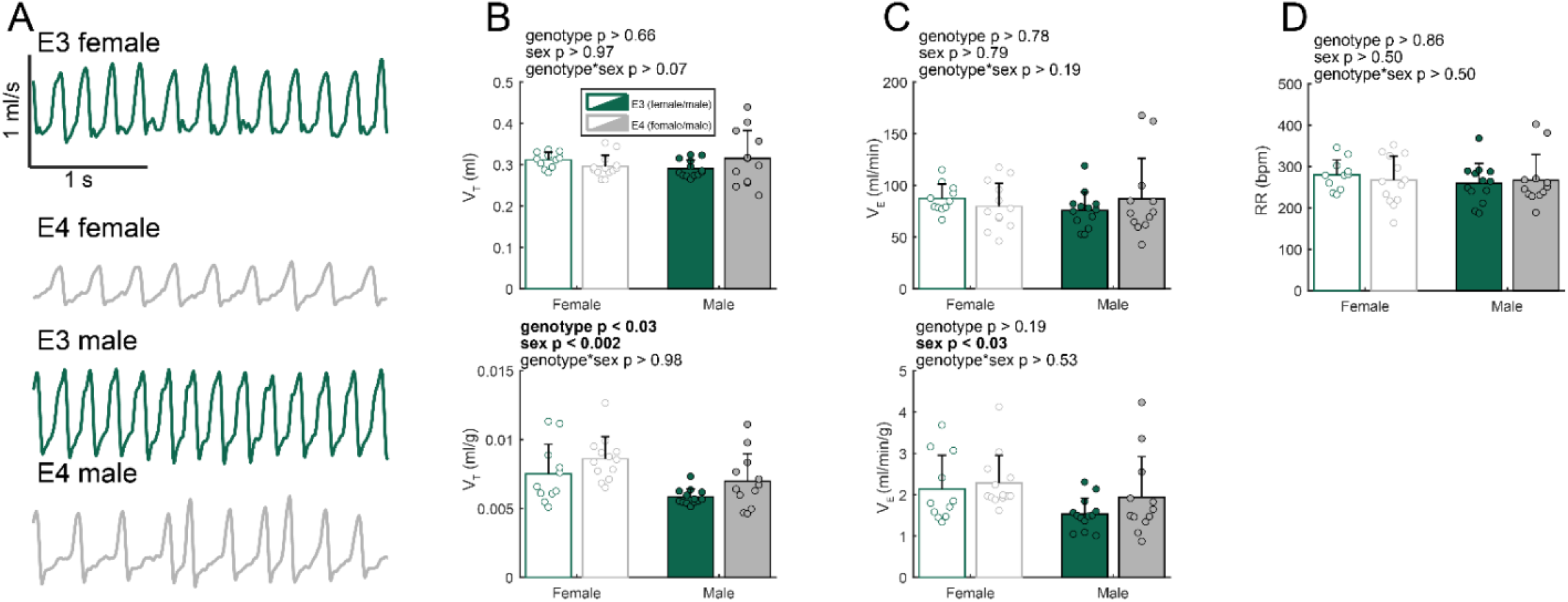
Female *APOE* mice breathe deeper than males to establish normoxic ventilatory patterns under stress-free conditions. **a)** Representative airflow traces during baseline (normoxia) from *APOE3* and *APOE4* male and female mice. Metrics of ventilation at baseline **b)** tidal volume (V_T_ (ml)) and weight corrected tidal volume (V_T_ (ml/g)), **c)** minute ventilation (V_E_ (ml)) and weight corrected minute ventilation (V_E_ (ml/g)), and **d)** respiratory rate (RR (bpm)). Male mice exhibit lower tidal volume and minute ventilation in weight corrected values, and *APOE4* animals exhibit greater tidal volume than *E3* counterparts across sexes. (Two-way ANOVA, mean + SD; n = 11 (*APOE3* female), n = 13 (*APOE4* female), n = 13 (*APOE3* male), n = 12 (*APOE4* male)).

### Male APOE3 mice exhibit a traditional HVR while female mice and APOE4 males fail to mount a complete response

The hypoxic ventilatory response is traditionally described by an increase in minute ventilation and tidal volume that peaks and is maintained for the duration of hypoxic challenge while frequency increases at first and followed by a slight decrease prior to leveling out at a rate higher than baseline frequency ^13^. To investigate sex- and genotype-dependent respiratory differences, we challenged the animals to a 10-minute hypoxic (11% O_2_) ventilatory challenge. Figure 3a shows airflow traces from animals undergoing hypoxic ventilatory challenge. Airflow data from all animals in the hypoxia cohort is quantified in Figure 3b, showing the hypoxic challenge period, split into two 5-minute bins. Within the first five minutes of hypoxia exposure, there is a clear effect of sex in respiratory rate and minute ventilation. During this initial period of hypoxia, female mice had a reduced respiratory rate relative to baseline and male mice had an elevated rate (p < 0.002, F_(1)_ = 12.82).

**Figure 3:**
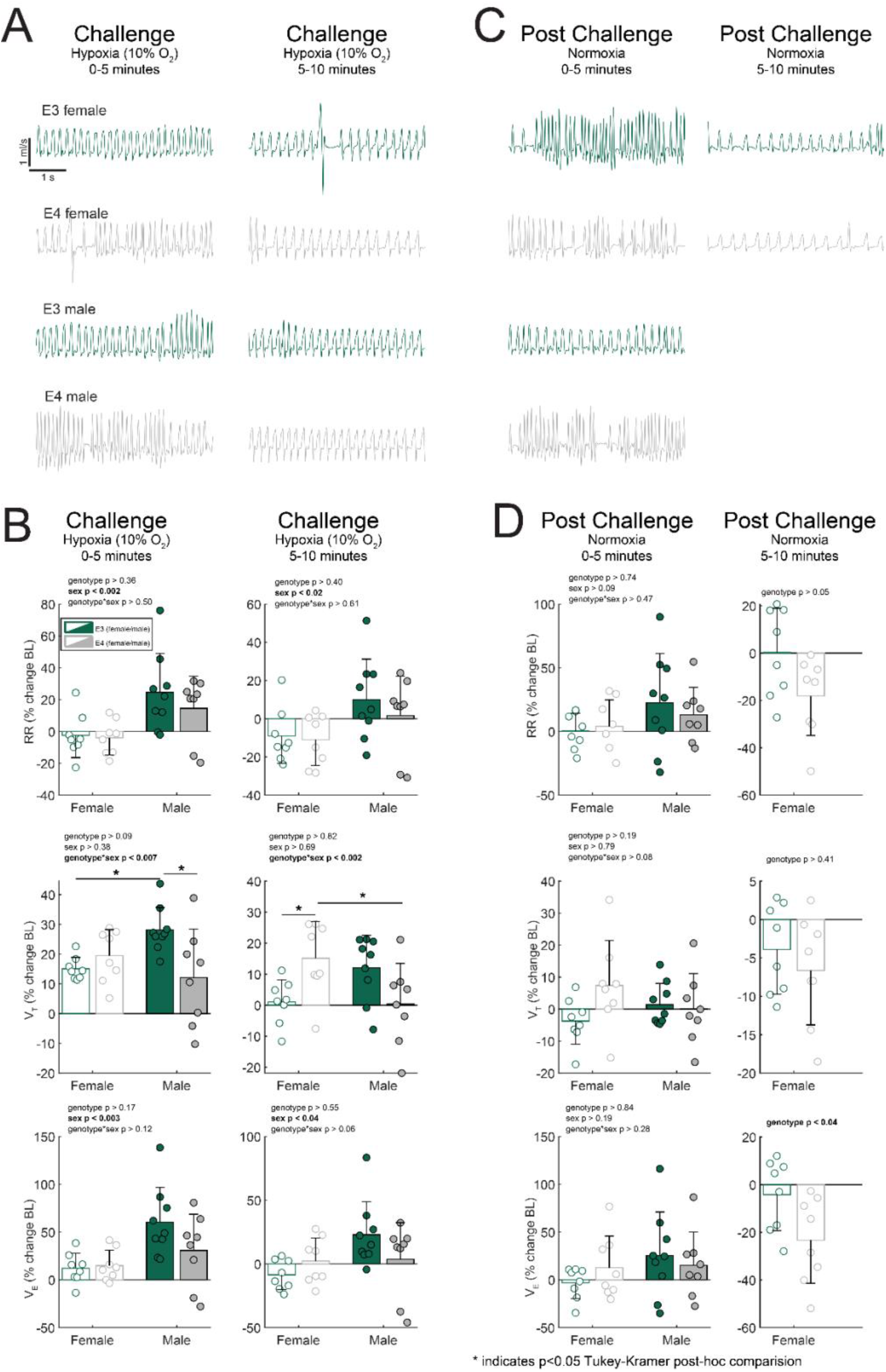
Male *APOE3* mice demonstrate a traditional HVR stronger than *APOE4* males and return to baseline levels following challenge while female mice fail to increase respiratory rate and *APOE4* females plummet to sub-baseline levels following challenge. **a)** Representative airflow traces during and after hypoxic ventilatory challenge. **b)** Comparison graphs between sex and genotype during hypoxic. **c)** Post-challenge airflow traces. **d)** Ventilatory metrics during the post-challenge normoxic period. Initial studies conducted in male mice did not include 5-10 minute post challenge period. Asterisks indicate p<0.05, Tukey-Kramer post-hoc. Two-way ANOVA, mean + SD; n = 8 (*APOE3* female), n = 8 (*APOE4* female), n = 9 (*APOE3* male), n = 8 (*APOE4* male).

During the final five minutes of the challenge, the respiratory rate of male mice returned towards baseline, while the respiratory rate of female mice declined further (p < 0.02, F_(1)_ = 6.57). All groups increased their tidal volume relative to baseline in the first five minutes of the hypoxic challenge. However, male *APOE4* mice had a less robust response compared to their *APOE3* counterparts (p < 0.05, Tukey-Kramer post-hoc), supporting previous work showing that male *APOE4* mice have impaired ventilatory response under hypoxic conditions after SCI ^14^. All groups increased their minute ventilation during the first 5 minutes, however, only the *APOE3* male mice were able to maintain an elevated minute ventilation over the entire challenge period. Interestingly, female mice remained at or below their baseline minute ventilation during the latter portion of the challenge while *E4* male mice could not maintain tidal volume increases in response to hypoxia.

### Response and recovery from hypoxia is impaired in female APOE3 and APOE4 mice

Post-hypoxia airflow traces show a return towards baseline level (Figure 3C). Following cessation of hypoxic challenge, there is a female genotype minute ventilation difference in the second 5-minute period (p<0.04 Figure 3D). When considering tidal and respiratory rate, female animals show a decline relative to baseline, with greater deficit seen in *APOE4* female mice. Interestingly, the female *E4* mice do have a slight initial tidal volume increase relative to baseline following the hypoxic challenge followed by a plummet below baseline levels and the levels of their female *E3* counterparts.

### During hypoxic hypercapnic challenge APOE4 animals are able to robustly increase respiratory output relative to APOE3 animals regardless of sex although sex plays a role during the post challenge period

As only the *APOE3* male group was able to produce and maintain a pronounced increase in minute ventilation during hypoxic challenge, we wanted to test if the other groups were limited in their ability to produce increased minute ventilation or if hypoxia alone simply did not trigger an increase. To accomplish this, in a separate cohort of animals, following the normoxic baseline period we challenged utilizing a hypoxic-hypercapnic (10.5% O_2_, 7% CO_2_) mixture. As the male *APOE4* animals during hypoxia did not increase their tidal volume to the same degree as the male *APOE3* animals, we were specifically interested in the possibility of a male *APOE4* ceiling effect (i.e., inability to generate larger tidal volume). Figure 4a shows a robust hypoxic-hypercapnic ventilatory response (HHVR) across all mice, regardless of sex or genotype. This suggests that there was not a ceiling effect within the *APOE4* male mouse hypoxia cohort and increases across all measures could be solicited via a more severe respiratory challenge indicative of impaired chemoreception in male *APOE4* mice.

**Figure 4:**
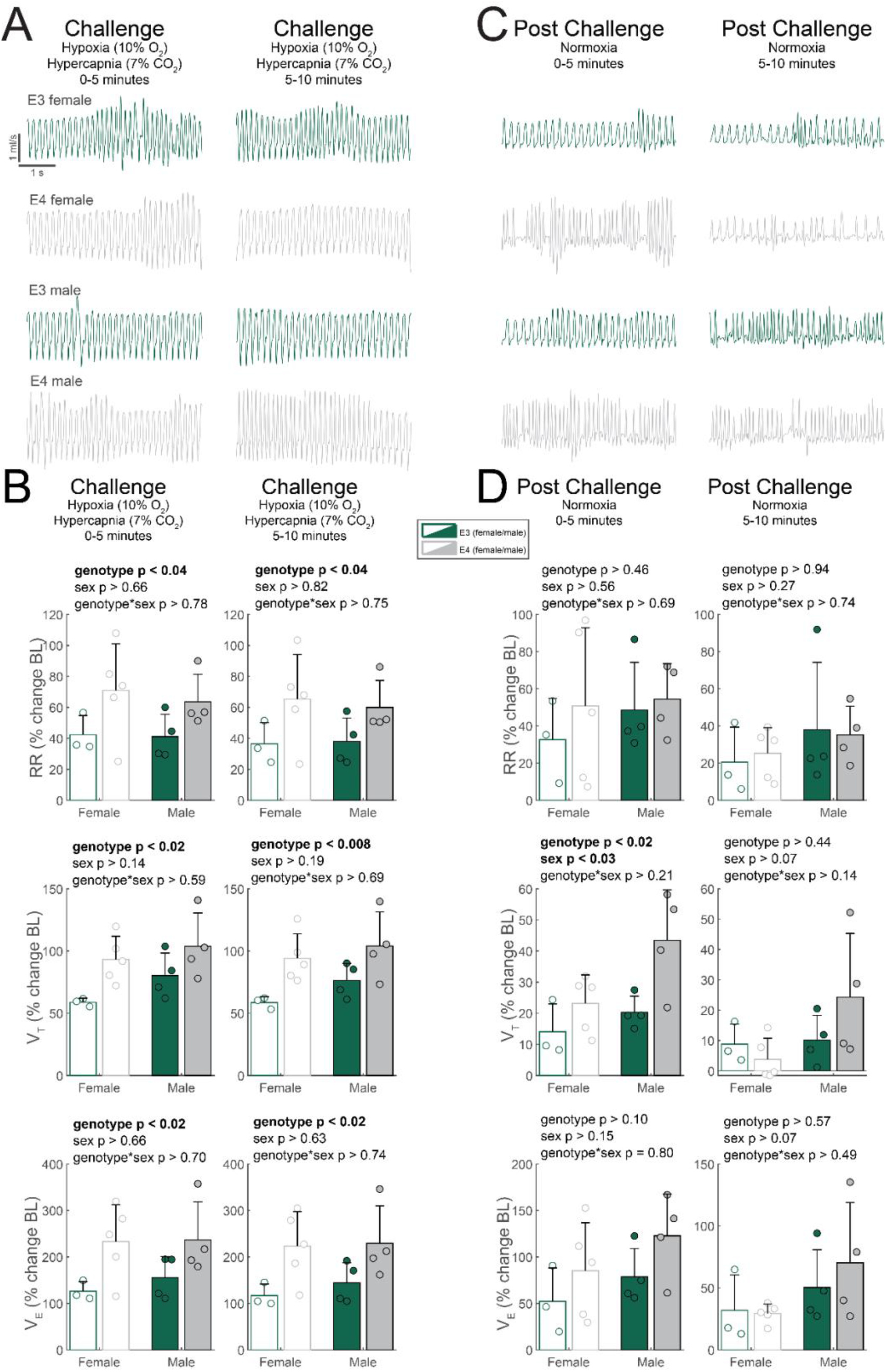
Under hypoxic hypercapnic conditions, *APOE4* animals respond robustly to challenge however *APOE4* males remain elevated relative to baseline following challenge. **a)** Flow traces show a robust increase in tidal volume during challenge. **b)** Comparison of ventilatory metrics during hypoxic-hypercapnic challenge, considering for effect of sex and genotype. **c)** Representative airflow traces during post-HHVR challenge. **d)** Respiratory metrics during the post-HHVR period. Two-way ANOVA, mean + SD; n = 3 (*APOE3* female), n = 5 (*APOE4* female), n = 4 (*APOE3* male), n = 4 (*APOE4* male).

Figure 4b shows the entirety of the hypoxic-hypercapnic challenge and the corresponding response. The *APOE3* mice of both sexes had a lower respiratory rate compared to *APOE4* counterparts during the first five minutes (p<0.04, F_(1)_ = 5.50) and the final five minutes (p<0.04, F_(1)_ = 5.61). Minute ventilation mirrors this effect throughout hypoxic-hypercapnic challenge, driven by genotypic differences in both respiratory rate and tidal volume, indicating that *E3* mice have lowered respiratory capacity relative to *E4* mice during hypoxic-hypercapnic challenge.

Airflow traces from the period after the hypoxic-hypercapnic challenge (Figure 4C) show a differential recovery based on sex and genotype. The male mice increased their respiratory rate in the first five-minutes after the challenge, while the female mice maintained the same respiratory rate as during the challenge. The tidal volume of all *E3* mice and *E4* female mice returned towards baseline, while the tidal volume of the *E4* male mice remained elevated.

The post-HHVR (Figure 4D) male mice have a higher tidal volume than female mice across both genotypes in the first (p<0.03, F_(1)_ = 6.93) 5 minutes with a trend in the second (p<0.08, F_(1)_ = 3.88) 5 minutes. This data may be indicative of impaired respiratory control in the male mice, as they should return toward baseline levels following exposure to hypoxic-hypercapnic conditions.

## DISCUSSION

This study builds on previous research elucidating the effects of genetic influences and sex on respiratory control ^14^. Breathing is a tightly regulated system that incorporates both feedback and feedforward control, however genetic disorders or predispositions can affect respiratory regulation ^3,4,5-7^. In prior work completed by our lab, female mice expressing the *E3* allele maintained diaphragmatic activity following intermittent hypoxia while *E4* females had significantly depressed activity. Unlike the female mice, males did not exhibit a difference in diaphragmatic activity based on genotype in our prior work. The results collected in this study continue to demonstrate that genotype and sex in humanized *APOE* mice plays a role in regulating breathing, specifically under conditions of respiratory stress. A summary of the individual changes can be referenced in Table 2.

**Table 2.** Summary of significant results across baseline measures and challenges.

| Normoxia | Baseline Measures |  |  |  |
| --- | --- | --- | --- | --- |
| Sex effects | Females have higher weight corrected **tidal volume**<br>Females have higher weight corrected **minute ventilation**. |  |  |  |
| Genotype effects | E4 animals have higher weight corrected **tidal volume**. |  |  |  |
| Interaction effects | none |  |  |  |
| Hypoxia | First 5 minutes | Second 5 minutes | Post-hypoxia first 5 minutes | Post-hypoxia second 5 minutes |
| Sex effects | Males have increased **respiratory rate**<br>Males have increased **minute ventilation** | Males have increased **respiratory rate**<br>Males have increased **minute ventilation** | none | none |
| Genotype effects | none | none | none | E4 females have depressed **minute ventilation** relative to E3 females |
| Interaction effects | E3 Males have increased tidal volume compared to **E3 females** and **E4 males** | E4 females have increased tidal volume compared to **E3 females** and **E4 males** | none | none |
| Hypoxic-Hypercapnia | First 5 minutes | Second 5 minutes | Post-hypoxia-hypercapnia first 5 minutes | Post-hypoxia-hypercapnia second 5 minutes |
| Sex effects | none | none | Males have increased **tidal volume** | none |
| Genotype effects | E4 animals have increased **tidal volume**, **respiratory rate** and **minute ventilation** | E4 animals have increased **tidal volume**, **respiratory rate** and **minute ventilation** | E4 animals have increased **tidal volume** | none |
| Interaction effects | none | none | none | none |

Severe OSA results in frequent periods of hypoxia which have been shown to increase neuroinflammation and impair cognition ^15,16,17^. In recent years, scientists have continued to address the impact of sex and age differences on OSA development ^18^. In addition to obstructive sleep apnea, approximately 1% of the United States population exhibits central sleep apnea resulting from impairments in central chemoreception ^19^. Severe respiratory impairments, including congenital central hypoventilation syndrome leading to fatal apneas, can result from genetic mutations ^20,21^. It follows that genetic polymorphisms may play a critical role in the onset and severity of OSA pathology and symptoms ^22,23^.

### Genotype and sex differences seen during hypoxia are reversed or eliminated when hypercapnia is introduced

While *APOE* genotype has been evaluated within sleep disordered breathing across broad datasets in human subjects, the research investigating animal models of *APOE* under respiratory stress has been limited ^24^. To model an adverse stimulus effect on respiratory pattern between *E3* and *E4* mice, we utilized hypoxic conditions eliciting the hypoxic ventilatory response investigating tidal volume, respiratory rate, and minute ventilation.

When exposed to hypoxia, our *APOE4* male mice were unable to maintain their increased frequency or tidal volume in response to this stimulus. These findings may have important implications in the context of obstructive sleep apnea for male *APOE4* carriers. While our *APOE4* female mice do generate a hypoxic ventilatory response specifically when considering tidal volume upon completion of the challenge they experience a massive drop in minute ventilation relative to *E3* female counterparts and well below their baseline minute ventilation. These changes point to either differences in respiratory responsiveness or reduced plasticity when encountering changes in pO2 which would result in an inability of *APOE4* carriers across sexes to adapt to periods of hypoxia during OSA ^25,26,27^. Indeed, the *E4* male carrier hypoxia response deficit may actually limit their ability to develop one possible compensatory mechanism of plasticity to overcome OSA impairments known as respiratory -long term facilitation (LTF) ^28^. This same interpretation can be applied to female *APOE4* mice who generate a robust HVR but at too great a cost.

While limited research asserts that *APOE4* may not contribute to OSA and vice versa our research has continued to show that the ability to attenuate a hypoxic insult such as that seen in OSA is linked to the *APOE4* genotype ^29,30^. As stated prior, hypoxia and the inability to compensate for it has been clearly shown to drive neuroinflammation and cognitive impairment. Chemoreceptor function under hypoxia is susceptible to genetic alterations demonstrated in carotid bodies, although models of humanized *APOE* mice have not yet been utilized ^31^.

Interestingly, our hypoxic hypercapnic ventilatory challenge data shows that *E4* animals can indeed increase their respiratory output in a stimulus dependent manner. During hypoxic hypercapnic challenge tidal volume and minute ventilation increased 50-200% relative to baseline. We show this across both genotypes and sexes with *E4* animals actually responding more robustly relative to *E3* counterparts. This is particularly important since it shows that *APOE4* animals have the ability to increase respiratory drive but not under conditions of hypoxia such as one would primarily experience during OSA. One reason for this may be that hypercapnic chemoreception is elicited primarily centrally in the retrotrapezoid nucleus while hypoxic chemoreception takes place peripherally in the carotid and aortic bodies ^32,33,34^. It could therefore be surmised that while mechanisms of peripheral chemoreception may be impaired in *E4* animals, central hypercapnic chemoreception remains unaffected or highly responsive to changes in pCO_2_ content. Since respiratory drive remains intact, pathways leaving brainstem respiratory centers may also be unaffected by *APOE*. These findings suggest that the elicitation of pCO2 chemoreceptive drive may represent one possible intervention for OSA individuals heterozygous or homozygous for *APOE4*.

Sex-dependent responses to hypoxia and hypoxic-hypercapnia have been studied in other animals, although the most relevant background to our current findings was a study in Fischer rats showing that ventilatory response differences are age-dependent as opposed to sex-dependent with middle aged and older female rats having a greater response ^35^. Sex differences in respiratory recovery have also been noted previously in neonatal mouse models where female respiratory rhythm generating centers have shown greater ability to recover following a period of severe hypoxia ^36^.

Future research will elucidate *APOE* specific differences in phrenic motor neuron serotonergic signaling ^14^. Additionally, future studies should include a cohort of aged matched wild-type male and female controls. The data presented here serves to fill a critical gap in knowledge as these *APOE* mice are both closer in relative age and more closely model conditions akin to human sleep apnea physiology ^37^.

## CONCLUSION

This study further clarifies work from our laboratory and others on the effects of *APOE* on respiratory control in unfavorable respiratory conditions. Our results show that humanized *APOE4* male mice have more difficulty mounting a response to hypoxic conditions than their *E3* counterparts while female mice of both genotypes appear to have difficulty mounting any response in terms of respiratory rate and only *E4* females maintain a respective tidal volume increase. Interestingly, *E4* female mice have prolonged depression following cessation of hypoxia, not seen in their male counterparts.

In hypoxic-hypercapnic conditions, genotype differences persist in across all measures, however, more interesting perhaps are the sex differences seen in recovery following hypoxic-hypercapnia. Because sleep apnea and disordered breathing have been indicated in earlier onset of disease as well as cognitive impairment this work will serve as a foundation for researching downstream effects of hypoxia and hypoxic-hypercapnia in genotype and sex allowing for the application of these findings to the heterogenous aging human population and to elucidating the development of disease ^38-42^.

## ACKNOWLEDGEMENTS

The study was supported by funding the Craig H. Neilsen Foundation (WJA) and the Kentucky Spinal Cord and Head Injury Research Trust (WJA).

Additional funding sources include NIH RO1s: AG065220 (LAJ), AG060056 (LAJ), AG80589 (LAJ), AG081421 (LAJ), AG066653 (RCS), CA266004 (RCS), AG078702 (RCS); the Alzheimer’s Association (LAJ), and the V-Scholar Grant (RCS).

## AUTHOR CONTRIBUTIONS

C.E.T.: design of experiment, data collection, data analysis and interpretation; initial draft of introduction and discussion. L.E.M.: data collection, data analysis and interpretation, initial draft of methods and results. M.D.S.: data analysis, statistical programming, final graphs, manuscript editing and revisions. J.N.W.: technical support, conceptual feedback. C.M.C.: technical support, conceptual feedback. R.C.S. data analysis and interpretation, manuscript editing and revisions: L.A.J.: mouse model breeding and support, review of manuscript draft. W.J.A.: conception and design of the study, experimental design, interpretation of the data, revision and final approval of the manuscript.

## DATA AND CODE AVAILABILITY

The datasets and MATLAB code generated during the current study are available from the corresponding author on reasonable request.

## ETHICAL DECLARATIONS

All experiments were conducted in accordance with the NIH Guidelines Concerning the Care and Use of Laboratory Animals and were approved by the IACUC committee at the University of Kentucky.

